# Restoration of β-globin expression with optimally designed lentiviral vector for β-thalassemia treatment in Chinese patients

**DOI:** 10.1101/2020.07.18.209759

**Authors:** Wenjie Ouyang, Guoyi Dong, Weihua Zhao, Jing Li, Ziheng Zhou, Gaohui Yang, Rongrong Liu, Yue Li, Qiaoxia Zhang, Xin Du, Haixi Sun, Ying Gu, Yongrong Lai, Sixi Liu, Chao Liu

**Affiliations:** BGI-Shenzhen, Shenzhen 518083, China; China National GeneBank, BGI-Shenzhen, Shenzhen 518120, China; Guangdong Provincial Key Laboratory of Genome Read and Write, BGI-Shenzhen, Shenzhen, 518120, China; Shenzhen Second People’s Hospital, First Affiliated Hospital of Shenzhen University, Shenzhen 518035, Guangdong, China; Department of Hematology, First Affiliated Hospital of Guangxi Medical University, Nanning, Guangxi 530021, China; Department of Hematology and Oncology, Shenzhen Children’s Hospital, Shenzhen, 518038, Guangdong, China

## Abstract

β-thalassemia is one of the most prevalent genetic diseases worldwide. The current treatment for β–thalassemia is allogeneic hematopoietic stem cell transplantation (HSCT), which is limited due to lack of matched donors. Gene therapy has been developed as an alternative therapeutic option for transfusion-ependent *β*-thalassemia (TDT). However, successful gene therapy for *β*-thalassemia patients in China has not been reported. Here we present the results of preclinical studies of an optimally designed LV named LentiHBB^T87Q^ in hematopoietic stem cells (HSCs) derived from Chinese TDT patients. LentiHBB^T87Q^ was selected from a series of LVs with optimized backbone and *de novo* cloning strategy. It contains an exogenous T87Q β-globin gene (HBB^T87Q^) driven by a specific reconstituted locus control region (rLCR) and efficiently express *HBB* mRNA and HBB protein in erythroblasts derived from cord blood (CB) HSCs. To facilitate clinical transformation, we manufactured clinical grade LentiHBB^T87Q^ (cLentiHBB^T87Q^) and optimized its transduction procedure. Importantly, transduction of cLentiHBB^T87Q^restored expression of HBB monomer and adult hemoglobin (HbA) tetramer to relatively normal level in erythroblasts from bone marrow (BM) HSCs of Chinese TDT patients, that carry the most common mutation types and cover various genotypes, including *β*0/*β*0. Furthermore, viral integration sites (VIS) of cLentiHBB^T87Q^ were similar to other LVs safely used in previous clinical trials and the associated risk of tumorigenesis was not observed in cLentiHBB^T87Q^ transduced HSCs through comprehensive analysis. Taken together, we have engineered the cLentiHBB^T87Q^ that can restore β-globin expression in the HSCs-derived erythroblasts of Chinese TDT patients with minimal risk on tumorigenesis, providing a favorable starting point for future clinical application.

## Introduction

*β*-thalassemia is the most prevalent hereditary blood disorder found in Mediterranean, Middle Eastern, Indian, South China and Southeast Asian^1, 2^. The patients in developing countries have shortened life expectancy and suffer from severe morbidity because of limited supportive therapies, imposing an enormous financial burden to the society and the patients’ families^3^. So far, the clinically available curative therapy for *β*-thalassemia is allogeneic hematopoietic stem cell transplantation (HSCT). However, for most patients, it is challenging to find the human leukocyte antigen (HLA) matched donors^4^. Recently, although the donor availability has been largely improved with the development of haploidentical HSCT (haplo-HSCT), patients receiving haplo-HSCT, particularly for those over 7 years old, still experience life-threatening complications due to graft versus host disease (GVHD)^5, 6^. Meanwhile, haplo-HSCT donors are usually patients’ parents with one mutated *β*-globin (HBB) allele showing a benign anemia phenotype, which may affect the patients’ final outcome and life quality following transplantation.

Gene therapy introduces functional HBB into patients’ own HSCs *ex vivo* and then infuse genetically corrected HSCs back through autologous transplantation. Since gene therapy uses patients’ own HSCs, donor availability and GVHD risk are largely avoided, providing an attractive therapeutic option^7–9^. Moreover, gene therapy with an efficient transduction and HBB expression, could theoretically restore HBB to a normal level and results in better clinical outcome when compared to most haplo-HSCT. Previous studies have shown successful correction of *β*-thalassemia in animal models using different lentiviral vectors (LVs)^10^. The first LV entering clinical trial was known as LentiGlobin, using a reconstituted local control region (rLCR) to express HBB with a T87Q mutation (HBB^T87Q^) that functions as wild type allele but meanwhile have anti-pathologic characteristic in sickle cell disease (SCD)^11, 12^. In phase I/II clinical trials, 12 of 13 patients with *β*0/*β*+ genotype and 3 of 9 patients with *β*0/*β*0 genotype became blood transfusion independent (TI) after gene therapy. The other 6 of *β*0/*β*0 patients showed a significant reduction (average ~73%) in annual transfusion volume. In consistent with the proposed mechanism, the overall clinical outcome was positively correlated to the inserted vector copy number (VCN) and exogenously expressed adult hemoglobin (HbA^T87Q^) characterized by a T87Q amino acid substitution^13^. Therefore, a higher VCN expressing sufficient HbA^T87Q^ is hypothesized to cure most *β*–thalassemia patients, including those with *β*0/*β*0 genotype. Indeed, in a latest phase III trial (NorthStar-III), with improved LV quality the mean VCN was increased to ~2.5 and the mean HbA^T87Q^ was above 9.5 mg/dL after gene therapy, enabling 75% of β0/ β0 patients to become TI for more than 3 months^14^. Up to date, LentiGlobin has treated more than 40 TDT patients and the earliest treated patients have been TI for over 36 months. Based on these promising data, gene therapy drug (Zynteglo) has been approved for treatment of non *β*0/*β*0 TDT patients over 12 years old in Europe^15^. However, the market price of this drug is 1.8 million dollars, which is too expensive for most TDT patients in developing countries. In another clinical trial, GLOBE LV drives exogenous HBB expression with a shorter LCR and, after gene therapy, 6 out of 7 treated patients decreased their blood transfusion volume dramatically, and 3 of them became TI for over one year. Interestingly, this study found that gene therapy was more effective in pediatric patients but didn’t increase the risk when compared to adult patients, suggesting it is ethical and reasonable to recruit pediatric and juvenile patients in future clinical trials^16^.

The aforementioned independent clinical trials have strongly suggested that gene therapy is promising for β-thalassemia treatment. Based on these clinical studies, the key to a successful gene therapy is to safely induce sufficient HBB expression, which is regulated by rLCR. The endogenous HBB LCR region is approximately 80kb in which the identified critical regions contain DnaseI hypersensitive sites (HSs), including HS1, HS2, HS3 and HS4, ranging from several hundreds of bp to over 2kb^17^. Different rLCRs have been constructed with various HS combinations to drive HBB expression^10, 18–21^. However, a systematic test to define the best rLCR combination with the highest efficacy has not been conducted. And finding a suitable, usually shorter rLCR, to facilitate the LV manufacture was not reported. Existing clinical used LVs including LentiGlobin and GLOBE are effective, but can’t help all TDT patients to get rid of transfusion, suggesting there is still room for improvement. Meanwhile, previous studies and clinical trials were mostly conducted in Caucasian and southeast Asian patients with genetic mutations of CD39, CD41/42, IVS-I-110 and HbE, etc^13, 16^. The most popular mutation types in south China including CD17, CD71/72 and IVS-II-654^22^ were not tested yet, and different genetic backgrounds among populations may be a complex factor for gene therapy. Therefore, we aim to develop an affordable gene therapy for Chinese TDT patients. Here, by optimizing rLCRs in the third-generation self-inactivating LV, we obtained and manufactured clinical grade LentiHBB^T87Q^ with robust HBB expression. With improved transduction procedures, we confirmed its efficacy in Chinese TDT patient samples and evaluated its safety profile regarding VIS, laying a foundation for further advancing this gene therapy to clinical studies in China.

## Materials and methods

### Lentiviral vector (LV) design, manufacture and titration measurement

The SIN vector of LentiHBB contains a CMV promoter and enhancer and deleted the U3 region of HIV in the 5’ LTR. The expression cassette is consisting of β-globin LCR and HBB mini-gene. LCR is composed of regions containing DNase I hypersensitive sites (HS), named as HS2, HS3 and HS4. HBB mini-gene comes with a short deletion of intron 2 and an anti-sickling amino acid substitution (T87Q). All of the DNA fragments were commercially synthesized (BEIJING LIUHE, Co. Ltd Guangzhou, China) and cloned into the LentiHBB vector using the Gibson assembly (NEB, E2611L). Lentivirus vectors of laboratory were produced by co-transfection of HEK293T cells with the transfer plasmid LentiHBB, the packaging plasmid psPAX2 and the envelope plasmid pMD.2G. The pseudotyped virions were collected from the clear supernatant and concentrated by ultracentrifugation at 80000g for 2h as previously described^23^. The clinical-grade LentiHBB^T87Q^ vector was manufactured according to good manufacturing practices (GMP) by OBiO Technology (Shanghai) Corp. Ltd.

The infectious titer of lentivirus was calculated by transducing HEK293T cells cultured in DMEM supplemented of 10% fetal bovine serum, glutamine, and polybrene (8 μg/mL). HEK293T cells was transduced with serial diluted lentivirus. The supernatant was replaced with fresh medium 24 hours after transduction and genomic DNA was extracted at the 3^rd^ day. VCN was measured by qPCR as described below. Titer was calculated multiplying the VCN/cell to the number of transduced cells and then normalized with dilution times, and was presented as transforming units per ml (TU/ml).

### HSCs culture and lentiviral transduction

Cord blood (CB) CD34+ cells were purified with Human Cord Blood CD34 Positive Selection Kit II (StemCell Technologies, 17896) and cultured in SFEM II medium (StemCell Technologies, 09605) supplemented with CD34+ Expansion Supplement (StemCell Technologies, 02691). Bone marrow (BM) and mobilized peripheral blood (mPB) CD34+ cells were purified with ClinMACS CD34 MicroBead Kit (Miltenyi,130-046-702) and cultured in in SCGM (CellGenix, 20802-0500) supplemented with 100ng/ml recombinant human cytokines thrombopoietin (TPO) (PrimeGene, 102-06), 100ng/ml fms-related tyrosine kinase 3 ligand (FltL) (PrimeGene, 103-05), 100ng/ml stem cell factor (SCF) (PrimeGene, 102-01) and 25ng/ml IL3 (PrimeGene, 101-03). CD34+ cells for transduction were cultured at a concentration of 1 × 10^6^ to 2× 10^6^ cells/ml at 37°C in a humidified incubator of 5% CO2. For lentiviral transduction, CD34+ cells were pre-stimulated in culture medium for 24 hours, and then transduced with lentivirus for 24 hours. The enhancers of protamine (Sigma, P4020), dmPGE2 (Cayman, 39746-25-3), UM171 (StemCell Technologies, 72914) and Poloxamer407 (BASF, 9003-11-06) were added at indicated concentration immediately prior to the addition of lentivirus.

### Methylcellulose Colony Forming Unit (CFU) assay

After lentiviral transduction, approximately 2000 HSCs were plated into Methocult Classic H4434 (StemCell Technologies, 04434) following the manual. After 14 days of culture, the number of Colony Forming Unit-Erythroid (CFU-E), Burst Forming Unit-Erythroid (BFU-E) and Colony Forming Unit-Granulocyte/Erythroid/Macrophage/Megakaryocyte (CFU-GEMM), Colony Forming Unit Granulocyte/Monocyte (CFU-GM) colonies were determined and scored based on colonies morphology, and individual BFU-E or pooled colonies were collected for VCN measurement and *HBB* expression analysis.

### Erythroid differentiation assay

CD34+ cells were washed with DPBS after lentiviral transduction or control treatment, and transferred to SFEMII medium (StemCell Technologies, 09605) supplemented with erythroid Expansion Supplement (StemCell Technologies, 02692). Erythroid differentiation medium was changed every 2 days and differentiated erythroblasts were collected for qPCR or HPLC analysis from day7 to day14.

### Genomic DNA (gDNA) extraction and Vector Copy Number (VCN) measurement

gDNA of LV transduced cells was extracted using the TIANamp Genomic DNA Kit (Tiangen, Beijin, China) following the manufacturer’s manual. gDNA of CFU colonies was extracted by suspending the cell in 1×PCR buffer (Takara, R050A) containing 0.2mg/ml proteinase K (Tiangen, RT403) and then incubating them at 55 °C for 20 min, 95 °C for 5min. The extracted genomic DNA can be stored at 4°C for short term usage or stored at −20°C for long term storage^24^.

VCN per diploid genome was determined by duplex TaqMan qPCR, using Premix Ex Taq kit (Takara, RR390A) and the StepOnePlus Real-Time PCR System (ABI, USA), following the manufacturers’ manuals. A plasmid (pMD18T-GAG-APOB) containing 1 copy of each GAG and APOB fragment was used to generate a standard curve, and copies of GAG and APOB were determined using this standard curve. The calculation of VCN (per diploid genome) was as follows^13^: VCN = [copies of GAG] ÷ [copies of APOB]×2.

qPCR Primers and probe sequences used for APOB and GAG were:

APOB qPCR F: 5-TGAAGGTGGAGGACATTCCTCTA-3;
APOB qPCR R: 5-CTGGAATTGCGATTTCTGGTAA-3;
APOB Probe: 5-VIC-CGAGAATCACCCTGCCAGACTTCCGT-3-TAMRA;
GAG qPCR F: 5-GGTTGTAGCTGTCCCAGTATTTGTC-3;
GAG qPCR R: 5-GGAGCTAGAACGATTCGCAGTTA-3;
GAG probe: 5-FAM-ACAGCCTTCTGATGTTTCTAACAGGCCAGG-3-TAMRA.

### Total RNA extraction and quantitative RT-PCR

Total RNA was extracted from HSC, erythroblasts or BFU-E using TRIzol reagent (Life Technology, CA, USA) following the manual instruction. RNA concentration and purity were determined by NanoDrop 2000 Spectrophotometer (Thermo Scientific, USA). DNA contamination was eliminated with gDNA Eraser (Takara, RR047A) and reverse transcription of mRNA was conducted using the PrimeScript™ RT reagent Kit (Takara, RR047Q) with oligo(dT) primers. Relative quantification PCR were performed using TB Green Premix Ex Taq kit (Takara, RR820B) and the StepOnePlus Real-Time PCR System (ABI, USA) following the manufacturers’ manuals. The expression of *HBB^WT^* and *HBB^T87Q^* were calculated according to the 2^-ΔΔCt^ method using *GAPDH* as the internal reference control as previous described^21^. Primers used for SYBR green qPCR were as follows. To measure *GAPDH* mRNA: *GAPDH* qPCR F: 5-ACCCACTCCTCCACCTTTGA-3; *GAPDH* qPCR R: 5-CTGTTGCTGTAGCCAAATTCGT-3; To measure total *HBB* mRNA: *HBB* qPCR F: 5-GAAGTCTGCCGTTACTGCCC-3; *HBB* qPCR R: 5-AGCCTTCACCTTAGGGTTGC-3; To measure relative expression percentage of *HBB* and *HBBT^87Q^*: *HBB^WT^* F: 5-TCAAGGGCACCTTTGCCACA-3; *HBB^T87Q^* F: 5-TCAAGGGCACCTTTGCCCAG-3 and *HBB^T87Q&WT^* R: 5-AATTCTTTGCCAAAGTGATGGG-3.

### High performance liquid chromatography (HPLC) analysis of globin and hemoglobin

Hemolysates were made by lysing 10^5^ erythroblasts or BFU-E cells in 50ul HPLC grade water for 10 minutes. After centrifugation at 13200rpm for 10min, 20ul of supernatant of hemolysates was used for HPLC analysis. The type of globin chains and hemoglobin tetramers were characterized using Aeris™ 3.6 μm WIDEPORE C4, 200 Å, 150 x 4.6 mm, LC Column (00F-4486-E0, Phenomenex) at 220nm and 100×4.6-mm cation exchange column (104CT0315, Phenomenex) at 418nm respectively on a HPLC machine (Agilent 1260 Infinity II). Globin samples were eluted with a gradient mixture of solution A (water/acetonitrile/trifluoroacetic acid, 700:300:0.7) and solution B (water/acetonitrile/trifluoroacetic acid, 450:550:0.5). Hemoglobin were eluted with a gradient mixture of Solution A (Bis-Tris 40mM, NaN3 2mM; pH adjusted at 6.5 with acetic acid) and solution B (Bis-Tris 40mM, NaN3 2mM, NaCl 200mM; pH adjusted at 6.8 with acetic acid) without degas program. Analysis and peak integration were performed using Open LAB CDS Chem station software (Agilent). Lysate of cord blood RBCs and adult peripheral blood RBCs from healthy donors were used as controls.

### Viral integration sites (VIS) Identification

The genomic sequences flanking VIS were identified by linear amplification mediated PCR (LAM-PCR) method and next generation sequencing (NGS)^25, 26^. Briefly, genomic DNA from cLentiHBB^T87Q^ transduced cells was fragmented by sonication using a Covaris E220 Ultrasonicator. The fragmented DNA was then subjected to end repair and 3’ adenylation, ligated to a custom adaptor include a 10 base pair random unique molecular identifier (UMI) and a specific sequencing adaptor for BGI-Seq 500 platform. The viral LTR-host genomic DNA junctions were amplified by a semi-nested PCR using LTR-specific primers and the adaptor-specific primers. The second-round adaptor-specific primer contains a unique 10-nucleotide barcode which indexes the amplification products. The PCR products were purified with AmpureXP beads, pooled in equimolar ratios, subjected to DNA nanoball (DNB) generation and then sequenced using the BGI-seq 500 platform.

### Bioinformatics analysis of VIS data

Raw reads were filtered by SOAPnuke^27^ with the parameters “-l 15 -q 0.2 -n 0.05”. Reads were aligned to the lentivirus sequences to remove vector backbone sequences, and then mapped to plasmid sequences to remove potential plasmid contamination. Finally, the remaining reads were aligned to the human reference genome (GRCh38.p12) using HISAT2^28^. Samtools^29^ was used to sort and index the alignment files for visualization using the Integrative Genomics Viewer tool^30^. The sorted binary sequence analysis file (BAM files) was also used to generate WIG files using BEDTools^31^. To this end, coverage files were normalized using the total signal for each sample. We generated a FASTA file of insertion sites from genome sequences (GRCh38.p12) using the getfasta function from BEDTools^31^ and used ggseqlogo^32^ R packages (version 0.1) to create 15 bp sequence logos (5 upstream and 10 downstream) from the FASTA files. Insertion sites and sequence information were used as input files for functional annotation using ANNOVAR^33^. We selected the insertion sites located in gene exons. GO enrichment analysis of the gene sets was then performed on each sample using the clusterProfiler R package^34,35^.

## Results

### Lentiviral vector bone selection and optimization

We selected the third-generation lentiviral vector (LV) backbone, as it will only be packaged into functional lentivirus with the help of its three assistant vectors and, thus, has safer characteristics^36^. To improve the lentiviral titers, we tested replacing its endogenous RSV promoter with CMV promoter^36, 37^. In agreement with previous studies, we found that LV containing CMV promoter always resulted in a higher tilter (Supplementary Figure S1A). In accordance, CMV version of LentiHBBs induced a higher expression level of beta-globin (HBB) when transduced into K562 cells (Supplementary Figure S1B). Thus, the LV backbone containing the CMV promoter was used in the following experiments (Figure 1A).

**Figure 1.**
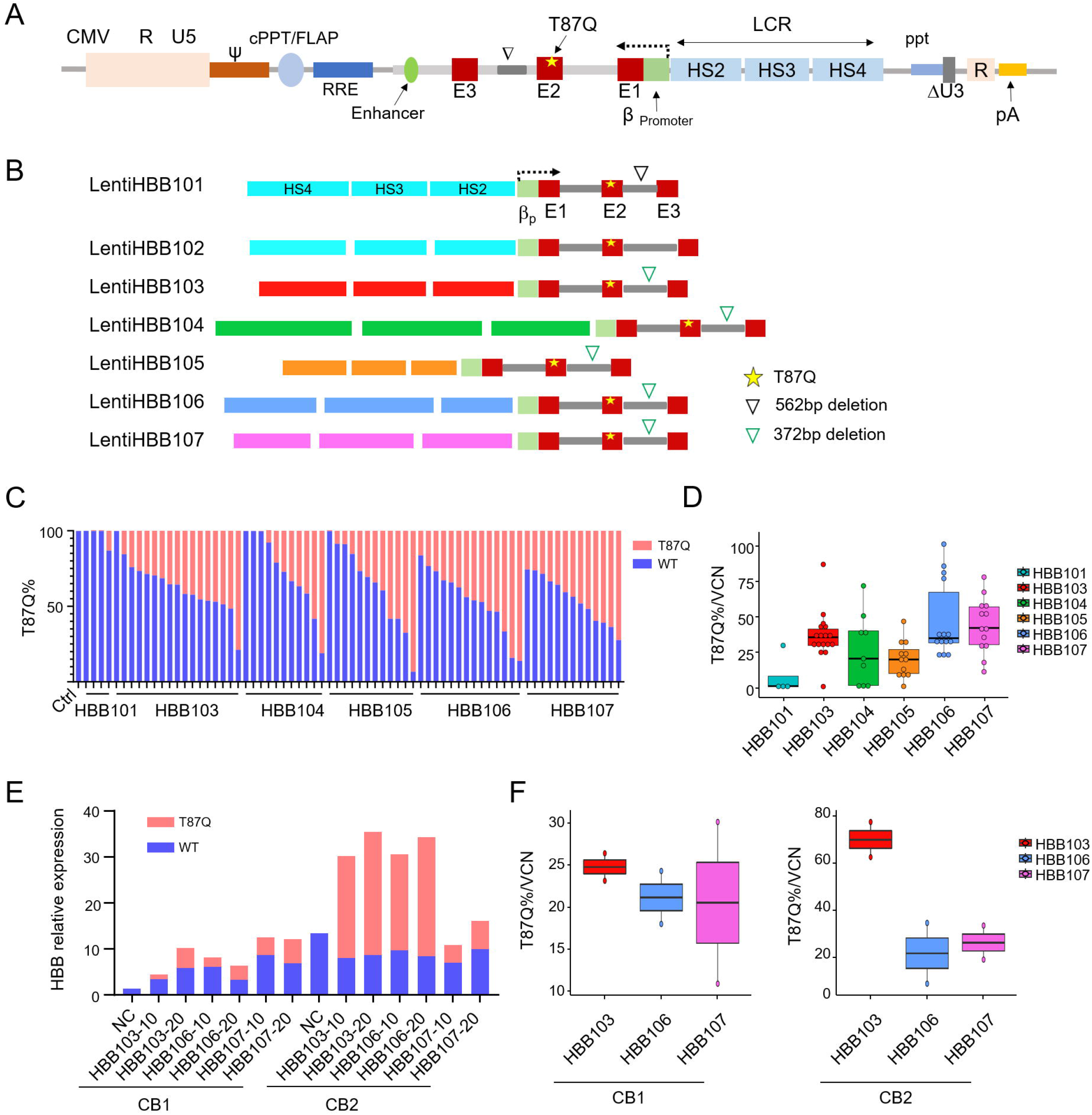
Construction and comparison of lentiviral vectors (LVs) containing different reconstituted local control regions (rLCR) in cord blood (CB) HSCs. (A) Diagram of our LV backbone, which contains a Cytomegalovirus (CMV) promoter and enhancer instead of the endogenous HIV promoter at the 5’ LTR, a deleted U3 region and a reversely inserted rLCR-HBB; (B) Diagram of different rLCR-HBBs constructed into LentiHBB101 to LentiHBB107; (C) Percentages of *HBB^T87Q^* mRNA in CFU-E colonies from CB HSCs transduced with different LVs; (D) Statistics of *HBB^T87Q^* versus VCN in CFU-E colonies shown in (C); (E) Percentages of *HBB^T87Q^* mRNA in differentiated erythroblasts from CB HSCs transduced with LentiHBB103, LentiHBB106 and LentiHBB107; (F) Statistics of *HBB^T87Q^* versus VCN in differentiated erythroblasts shown in (E). CB1 and CB2 are two independent samples.

### Construction of lentiviral vectors containing different reconstituted local control region (rLCR)

The local control region (LCR), critical for *HBB* regulation *in vivo*, contains 4 critical regions harboring DnaseI hypersensitivity site (HS). Prior studies using different reconstituted LCRs (rLCRs) with different combinations of HSs confirmed that most of them can drive *HBB* expression. However, no systemic comparison was performed between different rLCRs. Thus, we designed different rLCRs followed by HBB (rLCR-HBBs), cloned them into our CMV version of LV. To avoid blockade of lentiviral RNA transcription by HBB polyA, rLCR-HBBs were reversely inserted into the lentiviral backbone (Figure 1A). LentiHBB101 and LentiHBB102 contained the same 2.9kb rLCR^38^, and a 562bp deletion in the second intron of HBB was introduced in LentiHBB101 but not in LentiHBB102. LentiHBB103 to LentiHBB107 contained different rLCRs followed by the same HBB region that contains a 372bp deletion in the second intron (Figure 1B)^10, 18–21^. To avoid introducing other irrelevant sequences, which was the cases in LentiHBB101 and LentiHBB102, new rLCR-HBBs in LentiHBB103 to LentiHBB107 were synthesized *de novo* and then seamlessly cloned into LVs using Gibson Assembly^39^. Furthermore, the threonine at the 87th amino acid position was mutated to glutamine (T87Q) in HBB second exon. This mutated isoform (HBB^T87Q^) showed a stronger anti-sickling property and was used to treat sickle cell disease (SCD)^11, 12^. More important, *HBB^T87Q^* mRNA levels can be quantitively distinguished from the endogenous *HBB* mRNA using qPCR with sequence-specific probes.

### Expression of lentiviral vectors in erythroblast cells derived from cord blood HSCs

To measure exogenous *HBB ^T87Q^* expression levels, LentiHBBs were introduced to cord blood (CB) HSCs. Transduced HSCs were plated on methylcellulose to form single colonies, which were picked up for measurement of *HBB* expression and vector copy number (VCN). Percentages of each type of colonies were also calculated, and results indicated that transduction of different LentiHBBs did not affect the polyclonal ability of CB HSCs (supplementary Figure S2A). LentiHBB101 and LentiHBB102 produced a lentiviral tilter less than 1 × 10^8^ (Supplementary Figure S1A) and induced a negligible expression of exogenous *HBB ^T87Q^* in burst forming unit-erythroid (BFU-E) derived from the CB HSCs (Figure 1C, and data not shown). In contrast, LentiHBB103 to LentiHBB107 induced considerable *HBB ^T87Q^* expression in BFU-E and showed a high percentage of positive transduction. To compare their expression efficiency, we measured the exogenous *HBB^T87Q^* mRNA percentage (Figure 1C), and then normalized the *HBB^T87Q^* percentage to the inserted VCN in colonies of each group (Figure 1D). We found LentiHBB103, LentiHBB106 and LentiHBB107 were the top three LVs presenting efficient production of *HBB^T87Q^* mRNA, with LentiHBB103 showing the least variation (Figure 1D). We also found that LentiHBB103 had a higher correlation factor between *HBB^T87Q^* percentage and its VCN (Supplementary Figure. S2B), suggesting that LentiHBB103 documents consistent *HBB* expression.

To differentiate HSCs into erythroblasts, we cultured the transduced CB HSCs in erythroid medium for 2 weeks and then measured the expression levels of *HBB^T87Q^* mRNA in these pooled and differentiated cells. We found that the exogenous *HBB^T87Q^* mRNA expression level was sample dependent and positively correlated with the multiplicity of infection (MOI) (Figure 1E). Compared with LentiHBB106 and LentiHBB107, LentiHBB103 induced a higher expression levels of *HBB^T87Q^* mRNA after VCN normalization in two independent CB samples (Figure 1F). Thereafter, we chose LentiHBB103 to perform the following studies, and renamed it as LentiHBB^T87Q^. In summary, we generated a series of LVs containing different rLCRs, tested their *HBB^T87Q^* expression in erythroblasts derived from CB HSC samples, and identified the LentiHBB^T87Q^ with the highest efficacy and consistency.

### LentiHBB^T87Q^ express HBB protein in a dosage-dependent fashion

We further investigated whether the expressed *HBB* mRNA can be translated into HBB protein and form the hemoglobin tetramer in the differentiated erythroblasts. We transduced CB HSCs with LentiHBB^T87Q^ with MOI25, MOI50 and MOI100. Compared with the un-transduced control, the transduced samples showed an increased level of total *HBB* mRNA. VCN and *HBB^T87Q^* mRNA percentages were positively correlated with MOI values (Figure 2A). We also collected cell lysates for HPLC analysis of globin monomer and hemoglobin tetramer. In un-transduced control group, CB HSCs derived erythroblasts expressed high levels of α-globin and γ-globin, but negligible β-globin (Figure 2B). In agreement, the ratio of adult hemoglobin (HbA) to fetal hemoglobin (HbF) was as low as 0.16 in control group (Figure 2B-D). In contrast, the erythroblasts differentiated from the LentiHBB^T87Q^ transduced CB HSCs expressed a considerable amount of β-globin. When the MOI, and percentages of *HBB^T87Q^* mRNA was higher, the ratio of β-globin to α-globin was increased and the corresponding ratio of γ-globin to α-globin was decreased. In agreement, HbA to HbF ratio increased with the increase of MOI gradient and VCN (Figure 2B-D). The statistic and regression analysis showed *HBB^T87Q^* mRNA percentage and HbA/HbF ratio are positively correlated with VCN (Figure 2C, D). The high R^2^ value (above 0.9) indicates that lentiviral transduction mediated *HBB^T87Q^* expression increases HbA/HbF ratio. In conclusion, our studies show that LentiHBB^T87Q^ efficiently and dose-dependently generates HBB protein in erythroblasts derived from CB HSCs.

**Figure 2.**
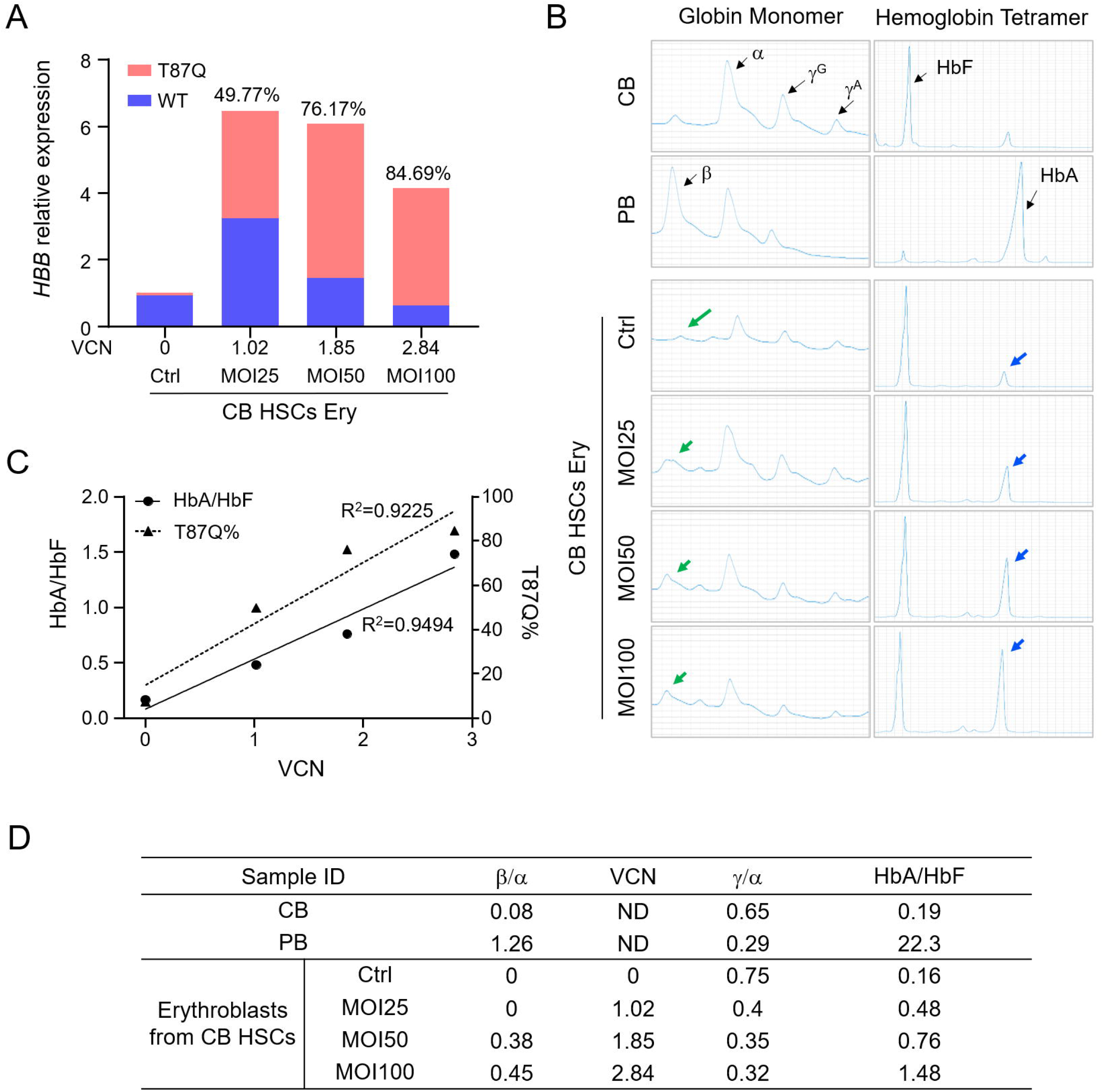
LentiHBB^T87Q^ increases *HBB^T87Q^* mRNA and HBB protein expression in erythroblasts derived from CB HSCs in a dosage dependent fashion. (A) The expression level of total *HBB* mRNA and the percentage of *HBB^T87Q^* in erythroblasts differentiated from CB HSCs transduced with LentiHBB^T87Q^; total *HBB* mRNA in transduced samples was normalized to control; (B) The globin monomer and hemoglobin tetramer of erythroblasts were determined by HPLC; Green arrows indicate the β-globin monomer and blue arrows indicate HbA tetramer; (C) Linear regression analysis among *HBB^T87Q^* percentage, HbA/HbF and VCN; (D) The statistic results of HPLC in panel B. ND, not determined.

### Optimization of lentiviral transduction in mobilized peripheral blood HSCs and validation of the optimized parameters with clinical-grade LentiHBB^T87Q^

For successful clinical application of gene therapy, it is critical to increase LV transduction efficiency while maintaining the cell stemness. Thus, it is important to optimize the *ex vivo* cell manipulation procedures, including the culture system, cell activation time, MOI value and the transduction enhancers, etc. As mobilized peripheral blood HSCs (mPB HSCs) is typically used for β-thalassemia gene therapy, we thus collected mPB HSCs from healthy donors for the optimization of the procedures for lentiviral transduction. First, we tested different combinations of cytokines, and determined their influence on transduction efficiency of LV and stemness of mPB HSCs. Among the tested three combinations, the one with the lowest content of cytokines showed the highest VCN and maintained the highest CD34+ percentage following LV transduction (Figure 3A). Second, we assessed distinct transduction enhancers, including protamine, dmPGE2, UM171, and Poloxamer407^40–42^ and tested their individual or combinational effects on LV transduction in mPB HSCs. We found that dmPGE2 and Poloxamer407 combination (PGE-Pol) was the most efficient, which increased the VCN by 6-7 times when compared with enhancer free control (Figure 3B). Meanwhile, PGE-Pol combination also showed the highest efficacy at different cell concentrations (Figure 3C) ranging from 1.0× 10^6^ to 2.0× 10^6^, suggesting that cell concentration had minimal effect on the transduction efficiency. Using the PGE-Pol enhancers, we transduced mPB HSCs with a gradient of MOI and measured the VCN and *HBB^T87Q^* mRNA percentage. The results indicated a positive correlation of VCN and *HBB^T87Q^* mRNA percentage with MOI (Figure 3D), and a positive correlation of *HBB^T87Q^* mRNA percentage with VCN in CFU-E (Figure 3E, F), suggesting that the usage of enhancers increased both VCN and the expression of *HBB^T87Q^* mRNA in mPB HSC samples. We also examined the cell activation time (from 16h to 48h) and LV transduction time (from 24h to 48h), and found no significant differences (data not shown). Last, we outsourced LV manufacture and obtained clinical grade LentiHBB^T87Q^, named as cLentiHBB^T87Q^. To validate the optimized procedures in clinical settings, we transduced mPB HSCs (~1.0×10^7^) that was purified with ClinMACS using cLentiHBB^T87Q^. The VCN up to 1.31 and *HBB^T87Q^* mRNA percentage up to 56.39% were positively correlated with MOI (Figure 3G). Although HPLC analysis cannot distinguish HbA^T87Q^ from WT HbA, we observed a clear shift of HbA position that was dependent on the increase of MOI. In contrast, HbF position was not affected by the transduction. Furthermore, HbA/HbF ratio was increased and positively correlated with MOI (Figure 3H, I), suggesting that increased expression of HbA was achieved by transduction of cLentiHBB^T87Q^. In addition, increased VCN upon increased MOI and positive correlation of *HBB^T87Q^* mRNA percentage with VCN were observed in CFU-E colonies from cLentiHBB^T87Q^ transduced groups (Figure 3J, K). Viability and stemness of mPB HSCs, as determined by FACS and CFU assays, were not altered by cLentiHBB^T87Q^ transduction (Figure 3L, M). Taken together, we have optimized the critical parameters of cell manipulation for *ex vivo* gene therapy and validated them with clinical grade LentiHBB^T87Q^ at a relatively large-scale in mPB HSCs, providing an entrance point for future therapeutic applications.

**Figure 3.**
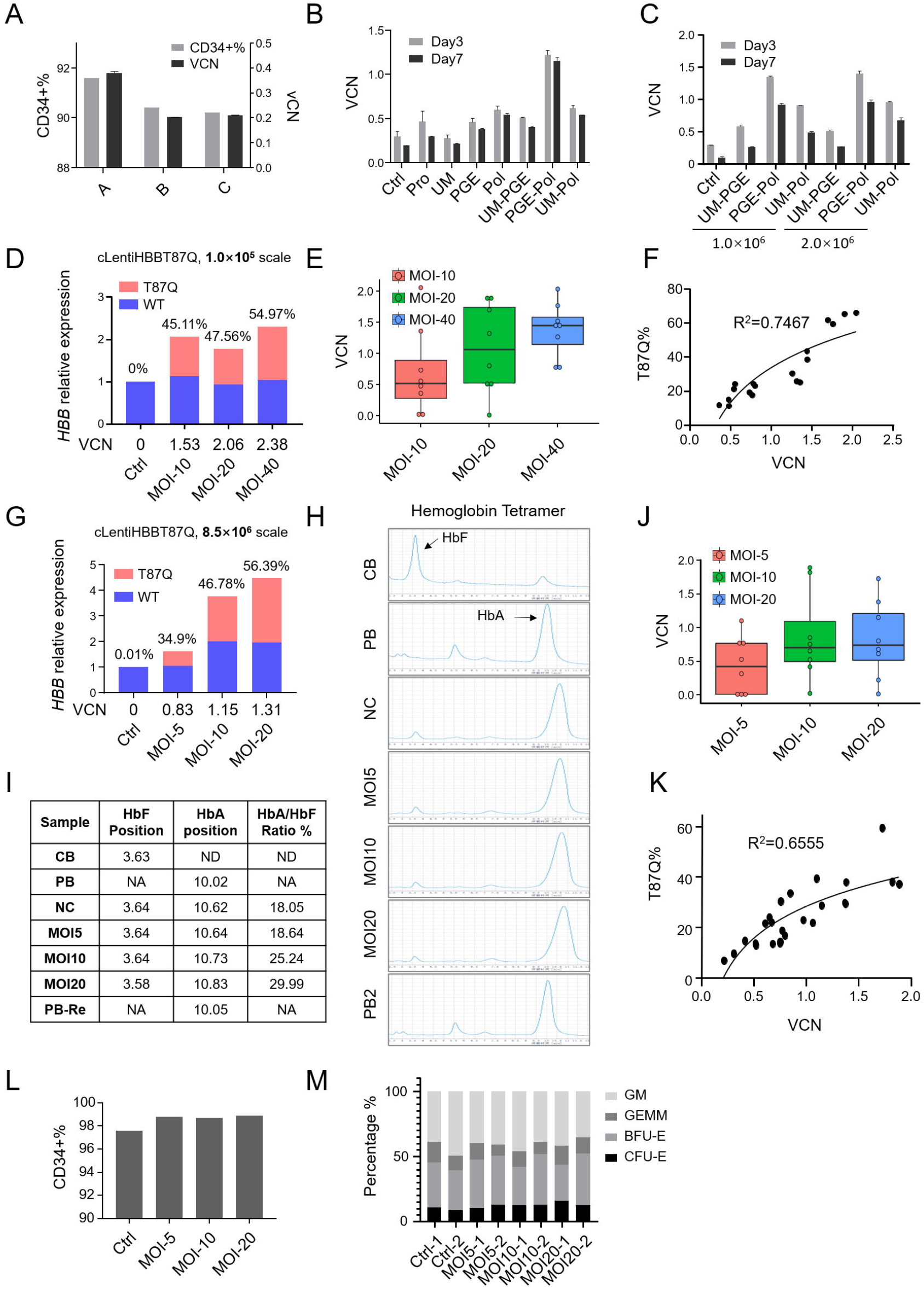
Optimization of lentiviral transduction in mobilized peripheral blood (mPB) HSCs and validation of the optimized parameters with clinical grade LentiHBB^T87Q^ (cLentiHBB^T87Q^) at different scales. (A) Results of CD34+ percentage and VCN after lentiviral transduction, when mPB HSCs were cultured in serum free medium with different cytokine combinations; (B) Results of VCN at 3 days and 7 days after lentiviral transduction when mPB HSCs were treated with different enhancer combinations during transduction. pro: protamine; UM: UM171; PGE: dmPGE2; pol: poloxamer407; (C) Results of enhancer combinations on transduction efficiency when mPB HSCs were cultured at different cell concentrations during transduction process; (D-F) Validation of transduction efficiency with clinical grade LentiHBB^T87Q^ (cLentiHBB^T87Q^) at 1.0 × 10^5^ cell scale; (D) VCN and *HBB^T87Q^* mRNA percentage were positively correlated with MOI in differentiated erythroblasts at 1.0 × 10^5^ cell scale; (E) Average VCN was positively correlated with MOI; (F) Linear regression of *HBB^T87Q^* percentage with VCN was obtained in CFU-E colonies that were derived from mPB HSCs transduced at 1.0× 10^5^ cell scale; (G-M) Validation of transduction efficiency with cLentiHBB^T87Q^ at 8.5 ×10^6^ cell scale; (G) VCN and *HBB^T87Q^* mRNA percentage were positively correlated with MOI in differentiated erythroblasts at 8.5 × 10^6^ cell scale; (H-I) HPLC and corresponsive statistic data of hemoglobin analysis in differentiated erythroblasts as shown in (G); ND, not determined; (J, K) Average VCN was positively correlated with MOI and linear regression of *HBB^T87Q^* percentage with VCN was observed in CFU-E colonies that were derived from mPB HSCs transduced at 8.5 ×10^6^ cell scale; (L, M) Results of CD34+ percentages and percentages of different colonies in CFU assay after transduction of cLentiHBB^T87Q^ at 8.5 ×10^6^ cell scale.

### Restoration of globin and hemoglobin with cLentiHBB^T87Q^ transduction in the erythroblasts differentiated from the bone marrow HSCs of Chinese TDT patients

To test the effect of cLentiHBB^T87Q^ on compensating *HBB* expression, we collected bone marrow (BM) samples from Chinese TDT patients (Figure 4A). Genotypes of these patients are β+/β0 and β0/β0 with the most common mutation alleles in South China. BM HSCs of TDT patients were purified and transduced with cLentiHBB^T87Q^ at MOI=20, following the procedures optimized in our aforementioned studies. All transduced samples have a VCN greater than 1.3 and showed increased total *HBB* mRNA levels, in which exogenous *HBB^T87Q^* mRNA amounted to more than 50% and up to 96.5% (Figure 4B). HPLC results indicated that expression of β-globin was profoundly increased and the ratios of β to α-globin were restored from less than 0.1 to approximately 1.0 in the β0/β0 samples transduced with cLentiHBB^T87Q^ (Figure 4C). In addition, erythroblasts from the transduced BM HSCs of TDT patients mainly expressed HbA^T87Q^ and largely eliminated the expression of HbF and other abnormal hemoglobin polymers that dominated in the un-transduced controls (Figure 4C). We also transduced a sample with cLentiBGI103 at lower MOIs of 5 and 10. The lower MOIs also resulted in efficient expression of *HBB^T87Q^* mRNA and HBB protein (Figure 4D), suggesting the high efficacy of cLentiHBB^T87Q^ and the robustness of our procedures. Taken together, our results suggest that cLentiHBB^T87Q^ can transduce HSCs from Chinese TDT patients *ex vivo*, and a VCN more than ~1.2 and a percentage of HBB^T87Q^ mRNA higher than ~50% were sufficient to produce significant improvements.

**Figure 4.**
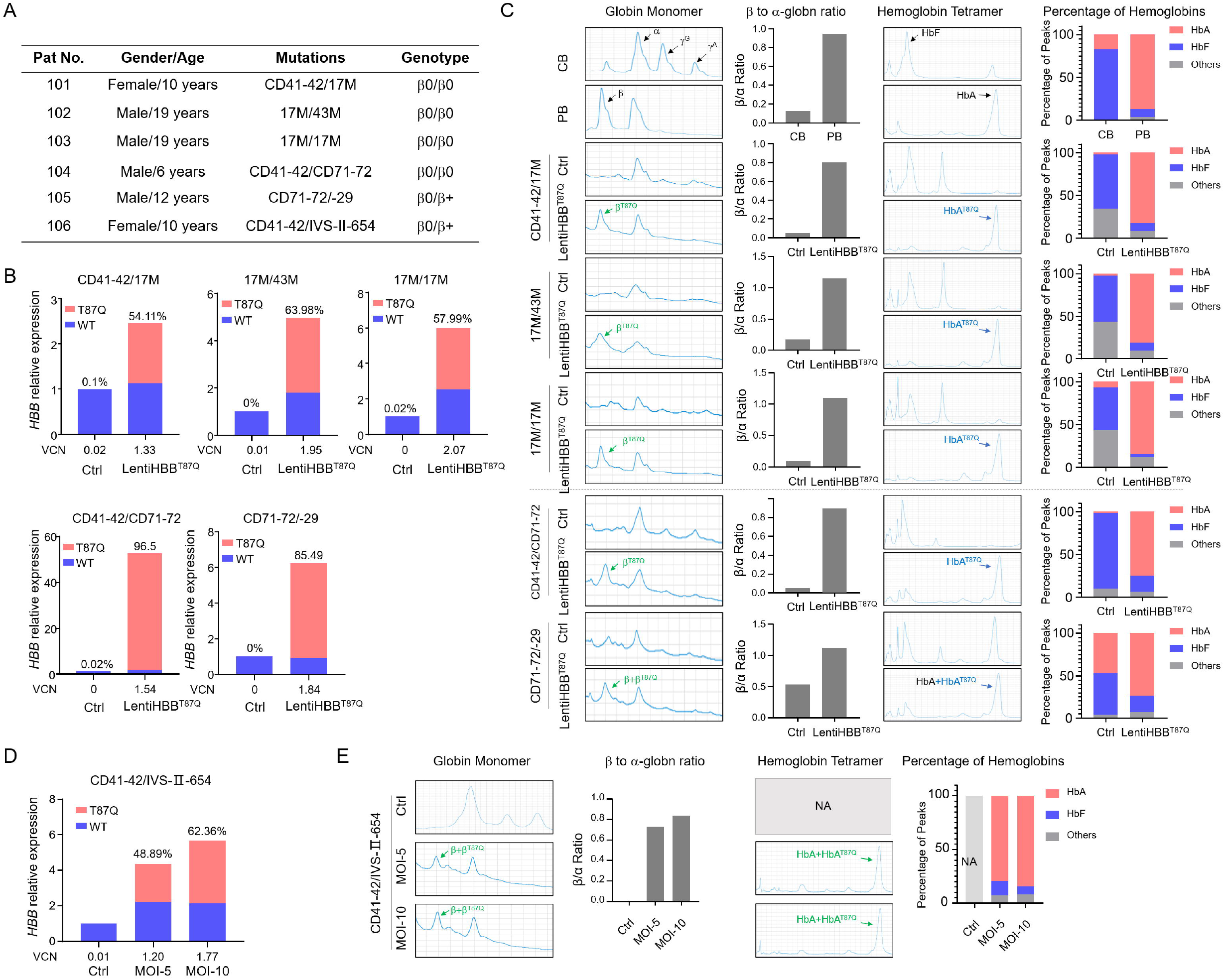
cLentiHBB^T87Q^ restores *HBB* mRNA and HBB protein expression in erythroblasts derived from Chinese TDT patients’ bone marrow (BM) HSCs. (A) Summary of the patients’ information; (B) Relative *HBB* mRNA expression level, VCN and *HBB^T87^* mRNA percentage after lentiviral transduction in erythroblasts derived from transduced BM HSCs of Chinese TDT patients; Genotype of each patient was list on top of the panel; (C) HPLC results of globin monomer and hemoglobin tetramer of erythroblasts from BM HSCs. The restored β^T87Q^ and HbA^T87Q^ were indicated by green and blue arrows respectively. Corresponsive statistic results of each HPLC panel were shown as columns on the right; (D) VCN and *HBB^T87Q^* mRNA percentage were positively correlated with MOI in TDT patients’ sample; (E) HPLC results of globin and hemoglobin in erythroblasts derived from patients’ HSC transduced with different MOI gradients.

### Characterization of viral integration sites (VIS)

To determine the safety spectrum of cLentiHBB^T87Q^, we analyzed the viral integration sites (VIS) in mPB HSCs transduced with cLentiHBB^T87Q^ at various MOIs (MOIs of 5, 10 and 20, as shown in Fig. 3G). The genomic DNA sequences flanking the VIS were retrieved by linear amplification mediated PCR (LAM-PCR) method (Figure 5A), sequenced using BGI next generation sequencing platform and mapped on the human genome (GRCh38.p12). Overall, we identified a large amount of unique VIS from transduced mPB HSCs. The VIS numbers ranged from 20885 to 38395, and were positively associated with MOI. Less than 5% overlapping sites were detected in different samples, suggesting that the insertion of cLentiHBB^T87Q^ were largely random (Figure 5B, C). The top10 most prevalent VIS account for less than 15% of total VIS, and contains two popular sites which were also detected as the top 2 sites in a previous clinical study (Figure 5D)^43^. VIS distribution profiles at the chromosome level and centered on transcription starting site (TSS) showed that consistent preference of cLentiHBB^T87Q^ for integration in gene-dense regions but far from TSS (>5kb), comparable with the patterns observed in gene therapy treated patient samples of previous clinical studies (Figure 5E, 5F)^16, 43, 44^. Consensus sequences of preferred insertion sites of cLentiHBB^T87Q^ were consistent between groups (Figure 5G). Finally, Gene-Ontology (GO) enrichment analysis of the genes targeted by VIS for biological processes, cellular components and molecular functions demonstrated no enrichment of tumor-related pathways (Figure. 5H). Collectively, our data indicate that the VIS of cLentiHBB^T87Q^ is similar to those of lentiviruses safely used in previous clinical studies, and based on our bioinformatics analysis, it did not increase the risk of tumorigenesis.

**Figure 5.**
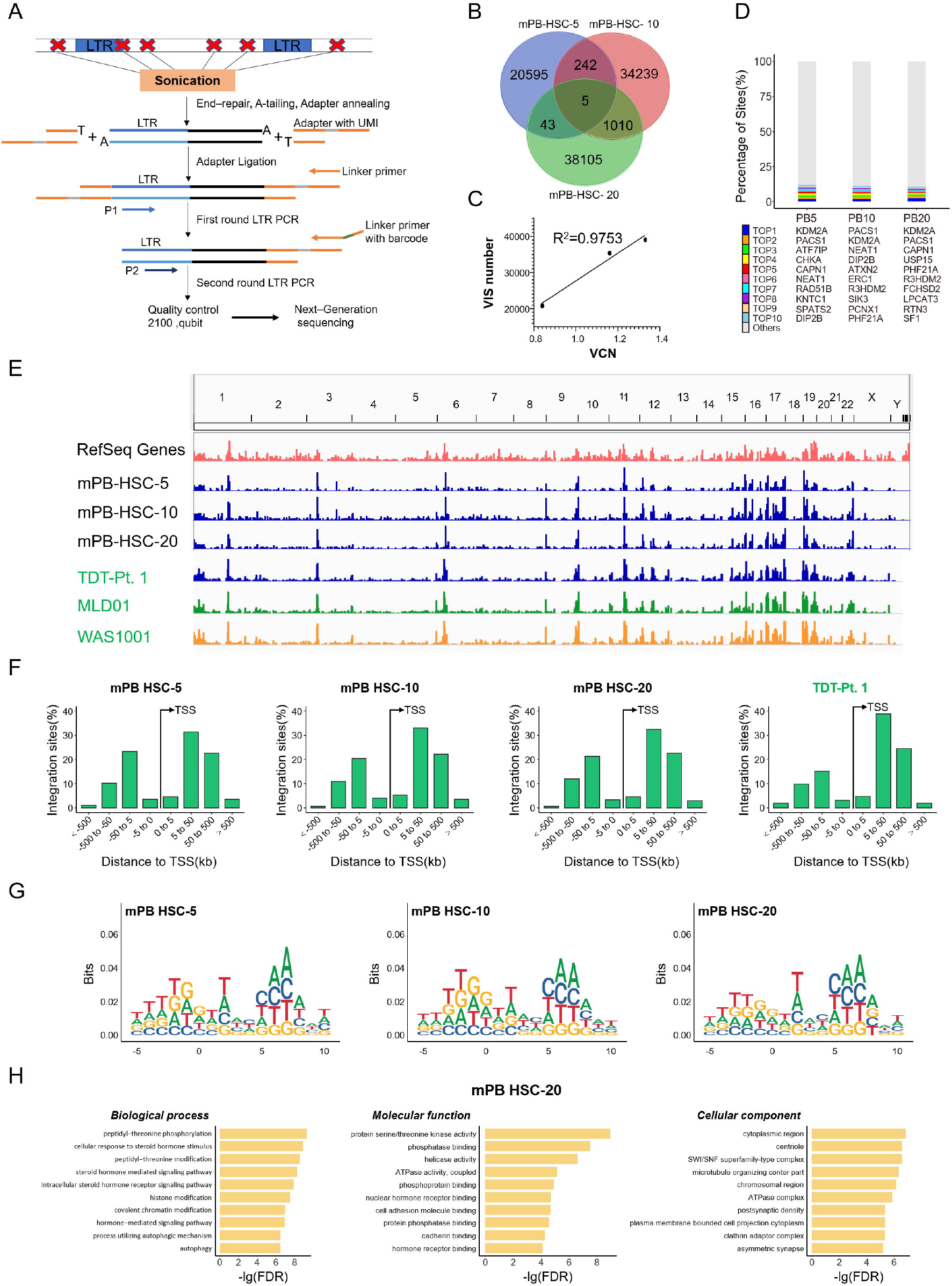
Characterization of viral integration sites (VIS) of cLentiHBB^T87Q^ transduced mPB HSCs. (A) Diagram of VIS identification process with linear amplification mediated PCR (LAM-PCR) and next-generation sequencing (NGS); (B) VIS numbers and overlaps among groups; (C) The VIS numbers is positively correlated with VCN; (D) Top ten integration sites in each group of cLentiHBB^T87Q^ transduced mPB HSCs; (E) Chromosome distribution profiles of VIS from samples transduced with cLentiHBB^T87Q^ and from patient samples transduced with other LVs in previous clinical trials; (F) Distribution of VIS centered on transcription starting site (TSS); Data cited from published papers were labelled in green in panel E and F; (G) The consensus sequences of preferred insertion sites of cLentiHBB^T87Q^; (H) Gene-ontology analysis of VIS targeted genes in a representative sample (mPB HSC-20) for biological processes, cellular components and molecular functions.

## Discussion

Gene therapy, as an attractive alternative option in addition to allogeneic HSCT, has been developed for β-thalassemia treatment. Two independent clinical studies indicated that the efficacy of gene therapy depends on exogenous *HBB* expression, which is driven by rLCR^13, 16^. Our results showed that rLCRs containing different Dnase I hypersensitive sites (HSs) drove *HBB* expression with different efficiency, and the rLCR with the longest HSs (LentiHBB104) did not necessarily deliver the best result (Figure 1D), suggesting that the core elements in HSs is sufficient to drive a high expression of *HBB* and other irrelevant sequences might play a negative role^21, 45^. Interestingly, the selected LVs containing the shorter rLCR named as LentiHBB^T87Q^ can be manufactured conveniently under GMP condition and achieve a high tilter up to 8.0 × 10^8^ TU/ml. The clinical grade LentiHBB^T87Q^ (cLentiHBB^T87Q^) transduced HSCs efficiently at a MOI as low as 5 (Figure 4D). Our data suggest an optimally designed cLCR not only increases the *HBB* expression efficacy but also facilitate the LV manufacture.

It is important to increase the transduction efficiency of HSCs *ex vivo* for a successful gene therapy. In our studies, we optimized the combinations of cytokines and transduction enhancers for cell culture and transduction. VCN in HSCs transduced with cLentiHBB^T87Q^ increased with the increase of MOI and reached up to ~2.38 (Figure 3D) at a small scale. However, mPB HSCs transduced with cLentiHBB^T87Q^ at a larger scale only had a VCN of 1.31 when MOI was 20 (Figure 3G). The lower VCN may be due to biological variation. Indeed, we noticed that VCNs were variable in the BM HSCs from different TDT patients, although a MOI was used at 20 in all samples. Nevertheless, VCN was always larger than 1.3 in all the transduced BM HSCs (Figure 4B). In a phase I/II clinical trial (HGB-205), cells with VCN above 0.6 produced HbA^T87Q^ level greater than 8 g/dL, and after infusion back to patients, dramatically alleviated thalassemia phenotypes^13^, suggesting that a VCN higher than 0.6 may be effective. In agreement, transduction of cLentiHBB^T87Q^ with our current procedures was sufficient to restore HBB and HbA expression *ex vivo* in all tested samples, including those with *β*0/*β*0 genotype (Figure 4). Interestingly, we found that increased abundance of HbF and other aberrant hemoglobins was present in the erythroblast differentiated from BM HSCs of TDT patients, but was largely eliminated after cLentiHBB^T87Q^ transduction, suggesting that exogenous *β*-globin^T87Q^ can competitively and stably bind to *α*-globin, in consistent with the fact that *β*-globin^T87Q^ has higher α-globin binding affinity and possesses anti-sickling function^46^. Therefore, like other LVs containing HBB^T87Q^, cLentiHBB^T87Q^ is expected to treat sickle cell disease (SCD) with improved potentcy^47^.

One potential risk for LV mediated gene therapies is tumorigenesis associated with viral integration, but the self-inactivation third generation of lentivirus has been used in over 100 clinical trials and shows a relatively safe profile^48^. Thousands of unique VIS of cLentiHBB^T87Q^ in transduced mPB HSCs were identified (Figure 5A-C). Consistent with prior studies^13, 16^, VISs are enriched in open chromatin regions (Figure 5E). Furthermore, VIS distribution and consensus sequence of the insertion locus show patterns comparable to those observed in previous clinical studies (Figure 5E-F), suggesting that cLentiHBB^T87Q^ may have similar characteristics to those clinically proven “safe” LVs. Our bioinformatics analysis of the genes influenced by VIS reveals no upregulation of cancer-related pathways, confirming that VIS of cLentiHBB^T87Q^ is unlikely to introduce malignant cellular transformation (Figure 5H). Future studies are required to monitor the resulting clonal expansion of cLentiHBB^T87Q^ transduced HSCs *in vivo*.

In summary, we have engineered a highly effective LV named as LentiHBB^T87Q^ for β-thalassemia treatment. We successfully manufactured clinical grade LentiHBB^T87Q^ (cLentiHBB^T87Q^) and show that the cLentiHBB^T87Q^ is robustly expressed in the mPB HSCs of healthy donors, and restores HBB function in the BM HSCs of Chinese TDT patients. Our systemic profiling of integration events in cLentiHBB^T87Q^ transduced HSC excludes the likelihood of increased tumorigenesis. Taken together, our data suggest cLentiHBB^T87Q^ is a promising candidate for the future therapeutic application in Chinese TDT patients.

## Data availability

The data that support the findings of this study have been deposited into CNGB Sequence Archive (CNSA: https://db.cngb.org/cnsa/) of CNGBdb with accession number CNP0001136.

## Acknowledgements

We thank members of BGI-research for helpful discussions; Dr. Longhou Fang for manuscript proof-reading and editing and Dr. Andrew Alpert for technical support on HPLC analysis. We sincerely thank the support provided by China National GeneBank.

## Ethics approval and consent to participate

Permission for this study was obtained from the Bioethics and Biological Safety Review Committee of BGI-Shenzhen (NO.16089-T2), the Medical Ethics Review Committee of Shenzhen Second People’s Hospital (NO. 20180515006), Shenzhen Children’s Hospital (NO.201900203) and First Affiliated Hospital of Guangxi Medical University (NO.2019001). We confirm that all experiments were performed in accordance with relevant guidelines and regulations.

## Author’s contribution

W.OY., Y.G., Y.R.L., S.L., and C.L. conceived and designed the experiments. W.OY., G.D., and J.L. performed the experiments. Z.Z and H.S. conducted the bioinformatics analysis. W.Z., G.Y., R.L., Y.L., Y.R.L. and S.L. provided the crucial clinical samples and collected the consent forms. Q.Z. and L.D. assisted in CD34+ cell purification. W.OY. and C.L. wrote the manuscript. All authors read and approved the final manuscript.

---

**Supplementary Figure 1. Efficacy comparison of CMV and RSV promoter in the 3rd lentiviral backbone.** (A) Tilter test of lentiviruses containing CMV or RSV promoter with rLCR-HBB ^Δ562^ or rLCR-HBB, and the CMV version of LentiHBB^Δ562^ and LentiHBB were LentiHBB101 and LentiHBB102 shown in Figure. 1B respectively; (B) Measurement of *HBB* mRNA induced by LVs containing CMV or RSV promoter with LCR-HBB ^Δ562^ or LCR-HBB in K562^-28 (G>A)^ mutant cell line, in which endogenous expression of *HBB* mRNA is eliminated by mutation of −28 site.

**Supplementary Figure 2. Comparison of different LentiHBB LVs containing different rLCR in cord blood (CB) HSCs.** (A) Statistic data of CFU colonies in each group after lentiviral transduction; (B) Nonlinear regression analysis of *HBB^T87Q^* mRNA with VCN in CFU-E from CB HSCs transduced with different LentiHBB.

## References

1. Modell B, Darlison M. Global epidemiology of haemoglobin disorders and derived service indicators. Bull World Health Organ 2008;86:480–487.

2. Piel FB, Patil AP, Howes RE et al. Global epidemiology of sickle haemoglobin in neonates: a contemporary geostatistical model-based map and population estimates. Lancet 2013;381:142–151.

3. Williams TN, Weatherall DJ. World distribution, population genetics, and health burden of the hemoglobinopathies. Cold Spring Harbor perspectives in medicine 2012;2:a011692.

4. Pennings G, Schots R, Liebaers I. Ethical considerations on preimplantation genetic diagnosis for HLA typing to match a future child as a donor of haematopoietic stem cells to a sibling. Hum Reprod 2002;17:534–538.

5. Faraci M, Bekassy AN, De Fazio V et al. Non-endocrine late complications in children after allogeneic haematopoietic SCT. Bone Marrow Transplant 2008;41 Suppl 2:S49–57.

6. Gaziev D, Galimberti M, Lucarelli G et al. Bone marrow transplantation from alternative donors for thalassemia: HLA-phenotypically identical relative and HLA-nonidentical sibling or parent transplants. Bone Marrow Transplant 2000;25:815–821.

7. Negre O, Eggimann AV, Beuzard Y et al. Gene Therapy of the β-Hemoglobinopathies by Lentiviral Transfer of the β(A(T87Q))-Globin Gene. Human gene therapy 2016;27:148–165.

8. Malik P. Gene Therapy for Hemoglobinopathies: Tremendous Successes and Remaining Caveats. Molecular therapy : the journal of the American Society of Gene Therapy 2016;24:668–670.

9. Mansilla-Soto J, Riviere I, Boulad F et al. Cell and Gene Therapy for the Beta-Thalassemias: Advances and Prospects. Human gene therapy 2016;27:295–304.

10. Negre O, Bartholomae C, Beuzard Y et al. Preclinical evaluation of efficacy and safety of an improved lentiviral vector for the treatment of beta-thalassemia and sickle cell disease. Current gene therapy 2015;15:64–81.

11. Takekoshi KJ, Oh YH, Westerman KW et al. Retroviral transfer of a human beta-globin/delta-globin hybrid gene linked to beta locus control region hypersensitive site 2 aimed at the gene therapy of sickle cell disease. Proceedings of the National Academy of Sciences of the United States of America 1995;92:3014–3018.

12. Pawliuk R, Westerman KA, Fabry ME et al. Correction of sickle cell disease in transgenic mouse models by gene therapy. Science 2001;294:2368–2371.

13. Thompson AA, Walters MC, Kwiatkowski J et al. Gene Therapy in Patients with Transfusion-Dependent beta-Thalassemia. The New England journal of medicine 2018;378:1479–1493.

14. Schneiderman JT, A.A.; Walters, M.C.; Kwiatkowski, J.L.; Kulozik, A.E; Sauer, M.G.; Porter, J.B; FRCP; FRCPath; Thuret, I.; Hongeng, S.; Lal, A.; Thrasher, A.J.; Yannaki, E; Elliot, H; Tao, G; Liu, WJ.; Colvin, R.A.; Locatelli, F.. Interim Results from the Phase 3 Hgb-207 (Northstar-2) and Hgb-212 (Northstar-3) Studies of Betibeglogene Autotemcel Gene Therapy (LentiGlobin) for the Treatment of Transfusion-Dependent β-Thalassemia. Biology of blood and marrow transplantation : journal of the American Society for Blood and Marrow Transplantation 2020.

15. Schuessler-Lenz M, Enzmann H, Vamvakas S. Regulators’ Advice Can Make a Difference: European Medicines Agency Approval of Zynteglo for Beta Thalassemia. Clinical pharmacology and therapeutics 2020;107:492–494.

16. Marktel S, Scaramuzza S, Cicalese MP et al. Intrabone hematopoietic stem cell gene therapy for adult and pediatric patients affected by transfusion-dependent ss-thalassemia. Nature medicine 2019;25:234–241.

17. Forrester WC, Takegawa S, Papayannopoulou T et al. Evidence for a locus activation region: the formation of developmentally stable hypersensitive sites in globin-expressing hybrids. Nucleic acids research 1987;15:10159–10177.

18. May C, Rivella S, Callegari J et al. Therapeutic haemoglobin synthesis in beta-thalassaemic mice expressing lentivirus-encoded human beta-globin. Nature 2000;406:82–86.

19. Roselli EA, Mezzadra R, Frittoli MC et al. Correction of beta-thalassemia major by gene transfer in haematopoietic progenitors of pediatric patients. EMBO molecular medicine 2010;2:315–328.

20. Breda L, Casu C, Gardenghi S et al. Therapeutic hemoglobin levels after gene transfer in beta - thalassemia mice and in hematopoietic cells of beta-thalassemia and sickle cells disease patients. PloS one 2012;7:e32345.

21. Weber L, Poletti V, Magrin E et al. An Optimized Lentiviral Vector Efficiently Corrects the Human Sickle Cell Disease Phenotype. Molecular therapy Methods & clinical development 2018;10:268–280.

22. Weatherall DJ. Phenotype-genotype relationships in monogenic disease: lessons from the thalassaemias. Nature reviews Genetics 2001;2:245–255.

23. Salmon P, Trono D. Production and titration of lentiviral vectors. Current protocols in human genetics 2007;Chapter 12:Unit 12 10.

24. Boulad F, Wang X, Qu J et al. Safe mobilization of CD34+ cells in adults with beta - thalassemia and validation of effective globin gene transfer for clinical investigation. Blood 2014;123:1483–1486.

25. Rae DT, Collins CP, Hocum JD et al. Modified Genomic Sequencing PCR Using the MiSeq Platform to Identify Retroviral Integration Sites. Human gene therapy methods 2015;26:221–227.

26. Schmidt M, Schwarzwaelder K, Bartholomae C et al. High-resolution insertion-site analysis by linear amplification-mediated PCR (LAM-PCR). Nature methods 2007;4:1051–1057.

27. Chen Y, Chen Y, Shi C et al. SOAPnuke: a MapReduce acceleration-supported software for integrated quality control and preprocessing of high-throughput sequencing data. Gigascience 2017;7:gix120.

28. Kim D, Langmead B, Salzberg SL. HISAT: a fast spliced aligner with low memory requirements. Nature methods 2015;12:357.

29. Li H, Handsaker B, Wysoker A et al. The sequence alignment/map format and SAMtools. Bioinformatics 2009;25:2078–2079.

30. Robinson JT, Thorvaldsdóttir H, Winckler W et al. Integrative genomics viewer. Nature biotechnology 2011;29:24.

31. Quinlan AR, Hall IM. BEDTools: a flexible suite of utilities for comparing genomic features. Bioinformatics 2010;26:841–842.

32. Wagih O. ggseqlogo: a versatile R package for drawing sequence logos. Bioinformatics 2017;33:3645–3647.

33. Wang K, Li M, Hakonarson H. ANNOVAR: functional annotation of genetic variants from high-throughput sequencing data. Nucleic acids research 2010;38:e164–e164.

34. Yu G, Wang L-G, Han Y et al. clusterProfiler: an R package for comparing biological themes among gene clusters. Omics: a journal of integrative biology 2012;16:284–287.

35. Ashburner M, Ball CA, Blake JA et al. Gene ontology: tool for the unification of biology. Nature genetics 2000;25:25.

36. Dull T, Zufferey R, Kelly M et al. A third-generation lentivirus vector with a conditional packaging system. Journal of virology 1998;72:8463–8471.

37. Miyoshi H, Blomer U, Takahashi M et al. Development of a self-inactivating lentivirus vector. Journal of virology 1998;72:8150–8157.

38. Ellis J, Pasceri P, Tan-Un KC et al. Evaluation of beta-globin gene therapy constructs in single copy transgenic mice. Nucleic acids research 1997;25:1296–1302.

39. Gibson DG, Young L, Chuang RY et al. Enzymatic assembly of DNA molecules up to several hundred kilobases. Nature methods 2009;6:343–345.

40. Heffner GC, Bonner M, Christiansen L et al. Prostaglandin E2 Increases Lentiviral Vector Transduction Efficiency of Adult Human Hematopoietic Stem and Progenitor Cells. Molecular therapy : the journal of the American Society of Gene Therapy 2018;26:320–328.

41. Uchida N, Nassehi T, Drysdale CM et al. High-Efficiency Lentiviral Transduction of Human CD34(+) Cells in High-Density Culture with Poloxamer and Prostaglandin E2. Molecular therapy Methods & clinical development 2019;13:187–196.

42. Ngom M, Imren S, Maetzig T et al. UM171 Enhances Lentiviral Gene Transfer and Recovery of Primitive Human Hematopoietic Cells. Molecular therapy Methods & clinical development 2018;10:156–164.

43. Biffi A, Montini E, Lorioli L et al. Lentiviral hematopoietic stem cell gene therapy benefits metachromatic leukodystrophy. Science 2013;341:1233158.

44. Aiuti A, Biasco L, Scaramuzza S et al. Lentiviral hematopoietic stem cell gene therapy in patients with Wiskott-Aldrich syndrome. Science 2013;341:1233151.

45. Leboulch P, Huang GM, Humphries RK et al. Mutagenesis of retroviral vectors transducing human beta-globin gene and beta-globin locus control region derivatives results in stable transmission of an active transcriptional structure. The EMBO journal 1994;13:3065–3076.

46. Nagel RL, Bookchin RM, Johnson J et al. Structural bases of the inhibitory effects of hemoglobin F and hemoglobin A2 on the polymerization of hemoglobin S. Proceedings of the National Academy of Sciences of the United States of America 1979;76:670–672.

47. Ribeil JA, Hacein-Bey-Abina S, Payen E et al. Gene Therapy in a Patient with Sickle Cell Disease. The New England journal of medicine 2017;376:848–855.

48. Milone MC, O’Doherty U. Clinical use of lentiviral vectors. Leukemia 2018;32:1529–1541.

