## Supplementary figures and images for "Restoration of β-globin expression with optimally designed lentiviral vector for β-thalassemia treatment in Chinese patients"

### OuyangSupplFig1.tif

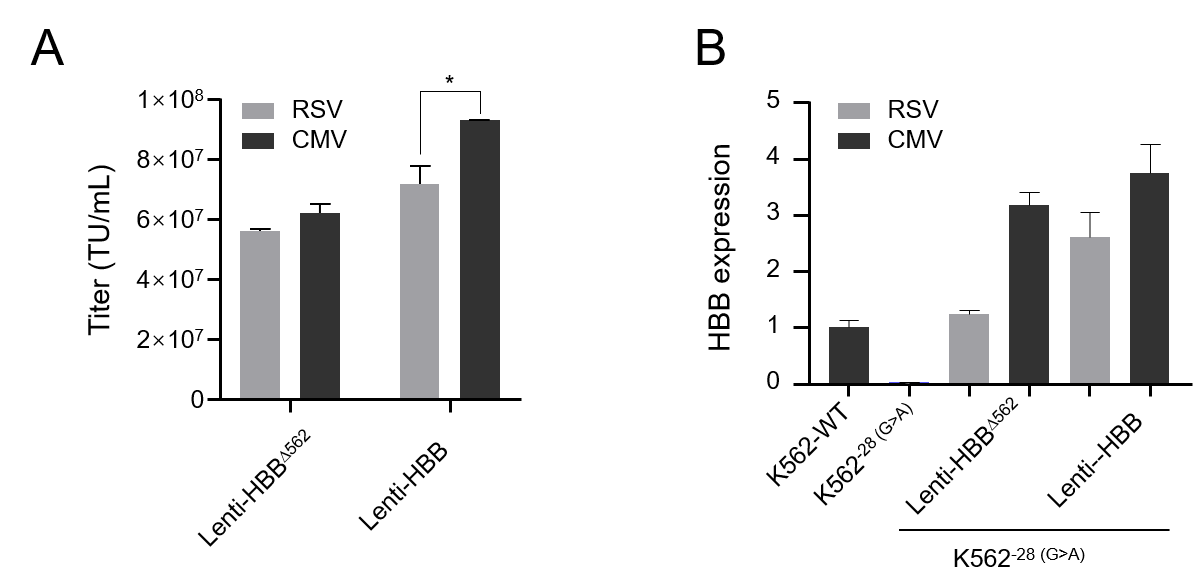

### OuyangSupplFig2.tif

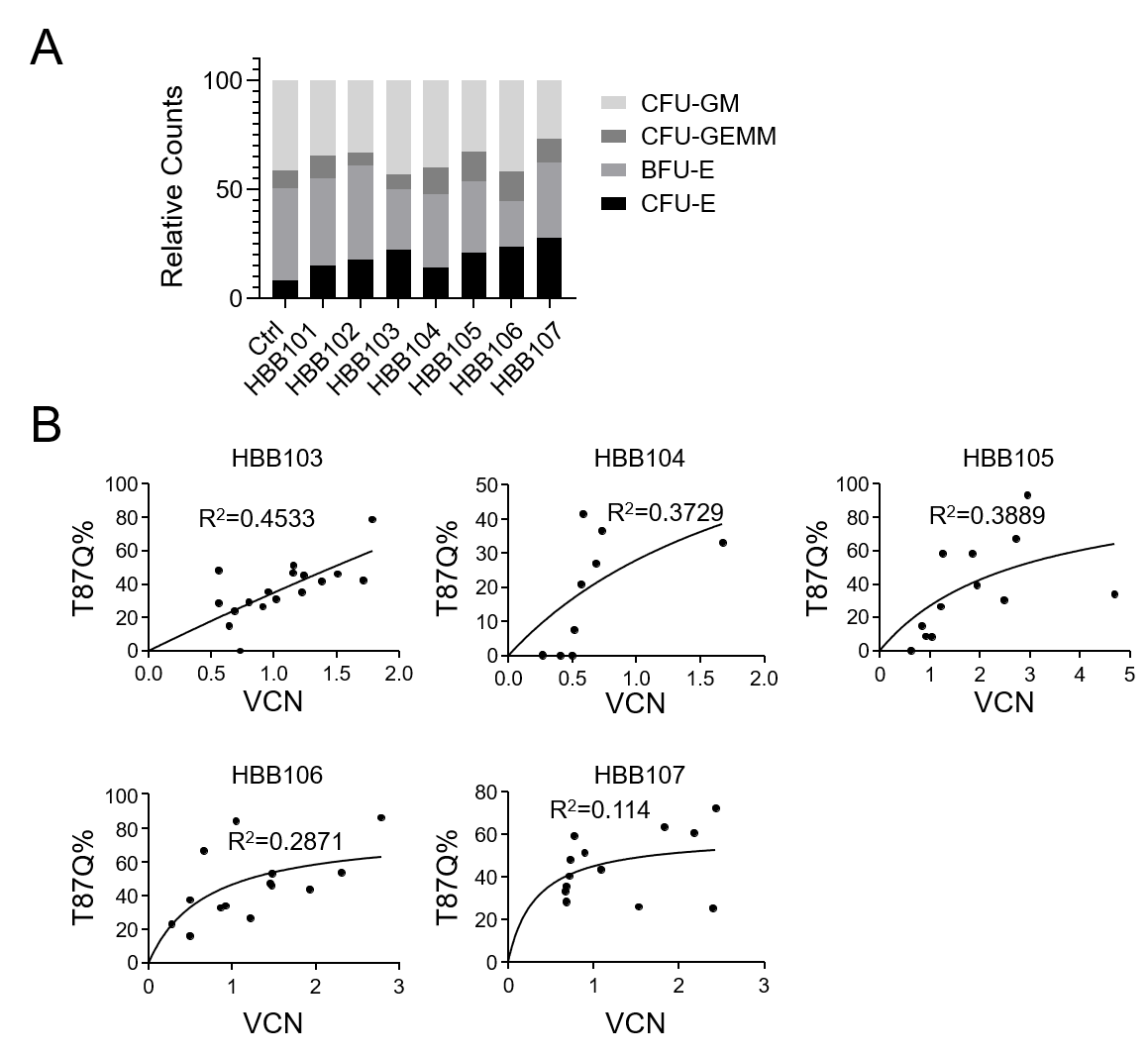
